# The utility of a metagenomics approach for marine biomonitoring

**DOI:** 10.1101/2020.03.16.993667

**Authors:** Gregory A. C. Singer, Shahrokh Shekarriz, Avery McCarthy, Nicole Fahner, Mehrdad Hajibabaei

## Abstract

The isolation and analysis of environmental DNA (eDNA) for ecosystem assessment and monitoring has become increasingly popular. A majority of studies have taken a metabarcoding approach—that is, amplifying and sequencing one or more gene targets of interest. Shotgun sequencing of eDNA—also called metagenomics—while popular in microbial community analysis has not seen much adoption for the analysis of other groups of organisms. Especially in light of the existence of extremely high-capacity DNA sequencers, we decided to test the performance of a shotgun approach side-by-side with a metabarcoding approach on marine water samples obtained from offshore Newfoundland. We found that metabarcoding remains the most efficient technique, but that metagenomics also has significant power to reveal biodiversity patterns, and in fact can be treated as an independent confirmation of ecological gradients. Moreover, we show that metagenomics can also be used to infer factors related to ecosystem health and function.

## 2 Introduction

The Earth’s oceans are under increasing environmental stress, and there have been calls for increased biomonitoring to understand these changes and (Canonico et al. 2019). Unfortunately, conventional observation-based techniques of biomonitoring are infeasible at such scales. However, the combination of DNA barcoding with next-generation DNA sequencing has led to the emergence of “metabarcoding” (Hajibabaei et al. 2011; Taberlet et al. 2012), and hundreds of studies over the past years have firmly established its utility as a means of rapid and accurate bioassessment.

Despite its capabilities, metabarcoding has some drawbacks. Although there exist a number of “universal” primers that purport to amplify common barcode regions over a very broad range of taxa, it is known that these primers don’t bind equally well to different templates, leading to biased amplification (Alberdi et al. 2018; Günther et al. 2018; Kelly, Shelton, and Gallego 2019). This is the primary reason why sequence read counts are only weakly correlated to the original starting DNA concentrations (Evans et al. 2016; Fonseca 2018; Piñol, Senar, and Symondson n.d.), and why abundance measurements from the technique are therefore unreliable. Moreover, these PCR biases can even lead to entire taxonomic groups missing (Tedersoo et al. 2015). Different techniques have been employed to ameliorate this bias, but it is most common to employ multiple gene regions (Porter and Hajibabaei 2018) and multiple primer pairs per gene target (Hajibabaei et al. 2019).

Alternatively, rather than amplifying and sequencing standardized DNA barcode regions from samples, it is possible to sequence random segments of DNA directly from the sample—essentially the whole genome shotgun sequencing of environmental samples, which is a technique commonly referred to as “metagenomics” (Handelsman et al. 1998; Chen and Pachter 2005; Tringe and Rubin 2005). Because there is no amplification of the DNA, the read counts should be directly proportional to the proportion of starting material, which itself is correlated to biomass (Yates, Fraser, and Derry 2019; Salter et al. 2019; Bista et al. n.d.). Curiously, although this was the first genomics approach used to characterize ocean biodiversity (Venter et al. 2004), it has since fallen out of favour except in the microbial sphere (Gregory et al. 2019) or in small systems such as the analysis of gut contents (Srivathsan et al. 2015) or small mock communities (Bista et al. n.d.). This is probably due to the relative inefficiency of the procedure: as shown in Table 1, while eukaryotic DNA is expected to be in low copy number within environmental samples (Azam and Malfatti 2007), the genome sizes are thousands of times larger than those of prokaryotes. Moreover, most of this extra DNA is noncoding, meaning it will not find a match in a reference database unless it happens to be closely related to an organism that has undergone whole genome sequencing. As has been observed by Stat et al (Stat et al. 2017), this leads to two outcomes: (1) the fraction of identifiable reads will be much lower than those from a metabarcoding experiment; and (2) while the reads may be an unbiased sample of the original eDNA, the *identifiable* reads will be highly biased towards genomes that have been completely sequenced—i.e., those from prokaryotes and eukaryotic model organisms. Indeed, the eukaryotic taxonomic recovery in previous studies is below 0.5% (Tedersoo et al. 2015; Stat et al. 2017).

**Table 1:** The likelihood of making a taxonomic classification on an environmental shotgun sequence read is an interplay between many factors, some of which are summarized here

| eDNA source | Abundance | Genome size | Reference database |
| --- | --- | --- | --- |
| Bacteria | ↑ | ↓ | ↑ |
| Eukaryotic nuclear | ↓ | ↑ | ↓ |
| Eukaryotic organellar | – | ↓ | ↑ |

With the recent introduction of the Illumina NovaSeq 6000—an ultra-high capacity DNA sequencing instrument that has a very low cost-per-read—we decided to revisit the utility of metagenomics for environmental characterization. We obtained samples from the near offshore of Newfoundland, Canada, and subjected those samples to both metabarcoding and metagenomics protocols.

## 3 Methods

### 3.1 Sample collection

Triplicate 250mL water samples were taken from surface water simultaneously. Samples were taken from eight locations along two transects in Conception Bay, Newfoundland and Labrador, Canada, on October 13-14, 2017 (Figure 1). These samples cover a range from nearshore to approximately 10km offshore (with a sea bottom depth ranging from a few metres nearshore to approximately 200 metres in the middle of the bay).

**Figure 1:**
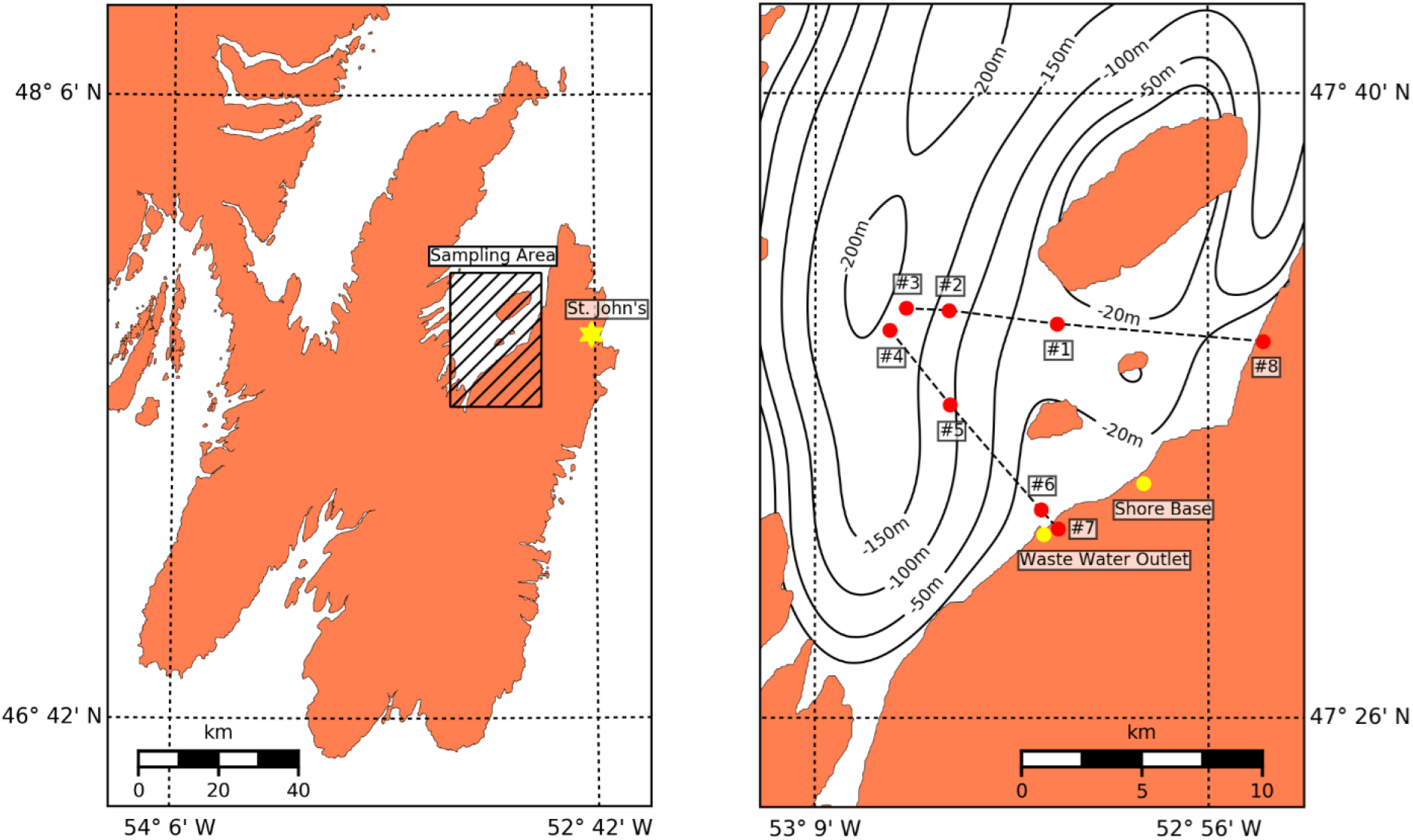
The location of eight sampling sites from Conception Bay, Newfoundland and Labrador, Canada.

### 3.2 Laboratory

Sample filtration, DNA extraction, and amplicon library prep were detailed in Singer et al. (2018). Briefly, water samples were filtered with 0.22 μm PVDF Sterivex filters (MilliporeSigma) and extracted with the DNeasy PowerWater Kit (Qiagen). Amplicons were then generated from the 232 bp Fish_miniE region of COI with triplicate PCR reactions (Shokralla et al. 2015). These amplicons were indexed with Nextera unique dual indexes (IDT). Amplicons were quantified with the Quant-iT PicoGreen dsDNA assay for normalization and library pooling. Libraries were cleaned with AMPure XP right and left side selections. Negative controls were generated during filtration, extraction, and PCR to screen for contamination and cross-contamination.

Metagenomic libraries were prepared with the Nextera Flex library prep kit (Illumina). For the majority of samples, 5 μL of DNA extract (1.0-3.0 ng/μL) was added to each tagmentation reaction. Three shore samples with higher DNA content were diluted so that 10 ng was added to each reaction (CBNL-052, CBNL-053, and CBNL-054, were diluted 32X, 4X, and 5X respectively). Ten PCR cycles were used to amplify and index tagmented DNA. Unique dual Nextera indexes were used to mitigate index misassignment (IDT; 8-bp index codes). Samples were quantified with Quant-iT PicoGreen dsDNA assay with a Synergy HTX plate fluorometer (BioTek) and pooled to normalize DNA concentration. Libraries were cleaned with three successive AMPure XP cleanups: Left side selection with bead:DNA ratios of 0.75X, then 0.8X, and a right-side selection with 0.5X.

Libraries were sequenced with a 300-cycle S4 kit on the NovaSeq 6000 following the NovaSeq XP workflow. Libraries were pooled with amplicon libraries from other projects for sequencing.

### 3.3 Bioinformatics

#### 3.3.1 Metabarcoding and metagenomics

All the raw amplicon reads as well as shotgun metagenomics reads, were trimmed for low-quality and Illumina sequencing adapters using Trimmomatic (Bolger, Lohse, and Usadel 2014). After filtering, there were a total of 132.8M total reads across the metabarcoding samples, and 217.2M reads across the metagenomics samples. A local k-mer based taxonomy database contains all of NCBI’s nt database constructed using Kraken (Wood and Salzberg 2014), and the taxonomic composition of both amplicon and shotgun reads predicted using Kraken. A threshold cut-off of 0.05 was used to report high confidence taxonomic assignments for each read. The taxonomic assignment tables were merged into one biom file format (McDonald et al. 2012) using an in-house python script and the data imported into the Phyloseq package in R (McMurdie and Holmes 2013) for downstream analysis. The relative abundance as well as read number tables constructed for each dataset. The Shannon metric values and Bray-Curtis distances were calculated for each sample using the Phyloseq package.

#### 3.3.2 Functional inference

Low-quality reads and Illumina universal adapters removed from the raw metagenomic reads using Trimmomatic (as above) and the quality of trimmed reads was assessed using fastqc (Andrews et al. 2012). The Comprehensive Antibiotic Resistance Database (CARD; Alcock et al. 2019) installed locally, and the shotgun reads per sample aligned against CARD’s protein homolog models using the Resistance Gene Identifier tool (Alcock et al. 2019). More specifically, FASTQ sequences aligned to the canonical CARD reference sequences that contain available sequences in GenBank with clear evidence for increased minimum inhibitory concentration (MIC) in a peer-reviewed journal available in PubMed.

## 4 Results and Discussion

### 4.1 Fewer metagenomics reads can be classified than metabarcoding reads

As noted above, the metabarcoding approach specifically targets genes that have large reference databases, whereas shotgun sequencing will produce random segments from nuclear and organellar genomes that may or may not have close matches in GenBank, and therefore it’s expected that a lower proportion of shotgun reads will be identifiable relative to metabarcoding reads. Indeed, this is exactly the case as illustrated in Figure 2. Regardless of the confidence threshold set for taxonomic classification, the result is the same: the majority of reads from metabarcoding studies can at least be identified to the level of kingdom, whereas the majority of reads from shotgun sequencing cannot even be identified at this very course level. The discrepancy is even more stark as attempts to map finer-level taxonomic information onto the reads are made. For example, while only 11.3% of metabarcoding reads were unclassified at the phylum level with 95% confidence, 62.7% of shotgun reads went unclassified.

**Figure 2:**
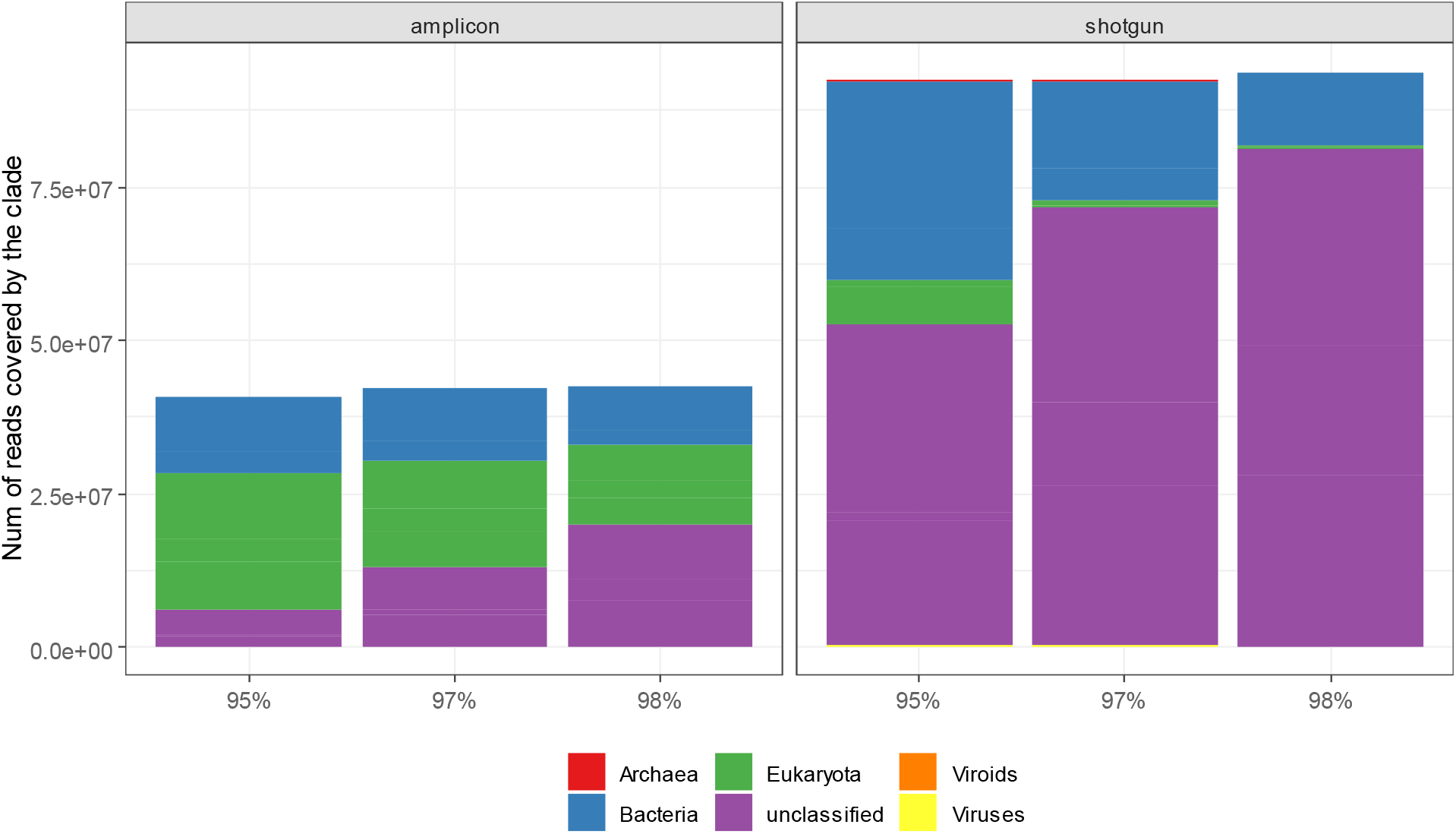
The proportion of unclassified reads (purple) is greater in the metagenomics experiment (right) than in the metabarcoding experiment (left). Data are plotted for a single sample (001). Note that the relative proportion of bacteria to eukaryotes among identified reads is much greater in the metagenomics experiment than in the metabarcoding experiment.

Figure 2 also upholds the second prediction made above: that taxonomic assignment to reads is also biased. Interestingly, we see opposite trends between the two methodologies here. In metabarcoding experiments, Eukaryota dominate the results—no doubt thanks to the enormous international efforts to build up these reference libraries (Adamowicz 2015). Conversely, the shotgun sequencing experiment shows a much larger proportion of Bacteria than Eukaryota—again likely the reflection of the relative completeness of the whole genome reference libraries to match against.

### 4.2 The pattern of diversity revealed by metagenomics is starkly different than that revealed through metabarcoding

If we restrict our analysis to just the Metazoan phyla, the two methodologies produce very different pictures of the biodiversity present in the environment (Figure 3). The metabarcoding experiments show a very high proportion of arthropods and few chordates, while this situation is reversed in the metagenomics experiment. This difference is even more stark when raw read counts are plotted (Figure 4). Note that despite the two-fold greater sequencing depth in the metagenomics experiments, the proportion of reads mapping to metazoan phyla is approximately half that of the metabarcoding experiments, again underscoring the relative inefficiency of the metagenomics approach. Also, the pattern of biodiversity is again starkly different—so much so that it is almost difficult to believe they came from the exact same DNA extracts. In this way, metagenomics data can be seen as complementary to metabarcoding data—both providing useful information that does not perfectly overlap. This is similar to the notion that metabarcoding data itself is complementary to traditional observational methodologies (Emilson et al. 2017)

**Figure 3:**
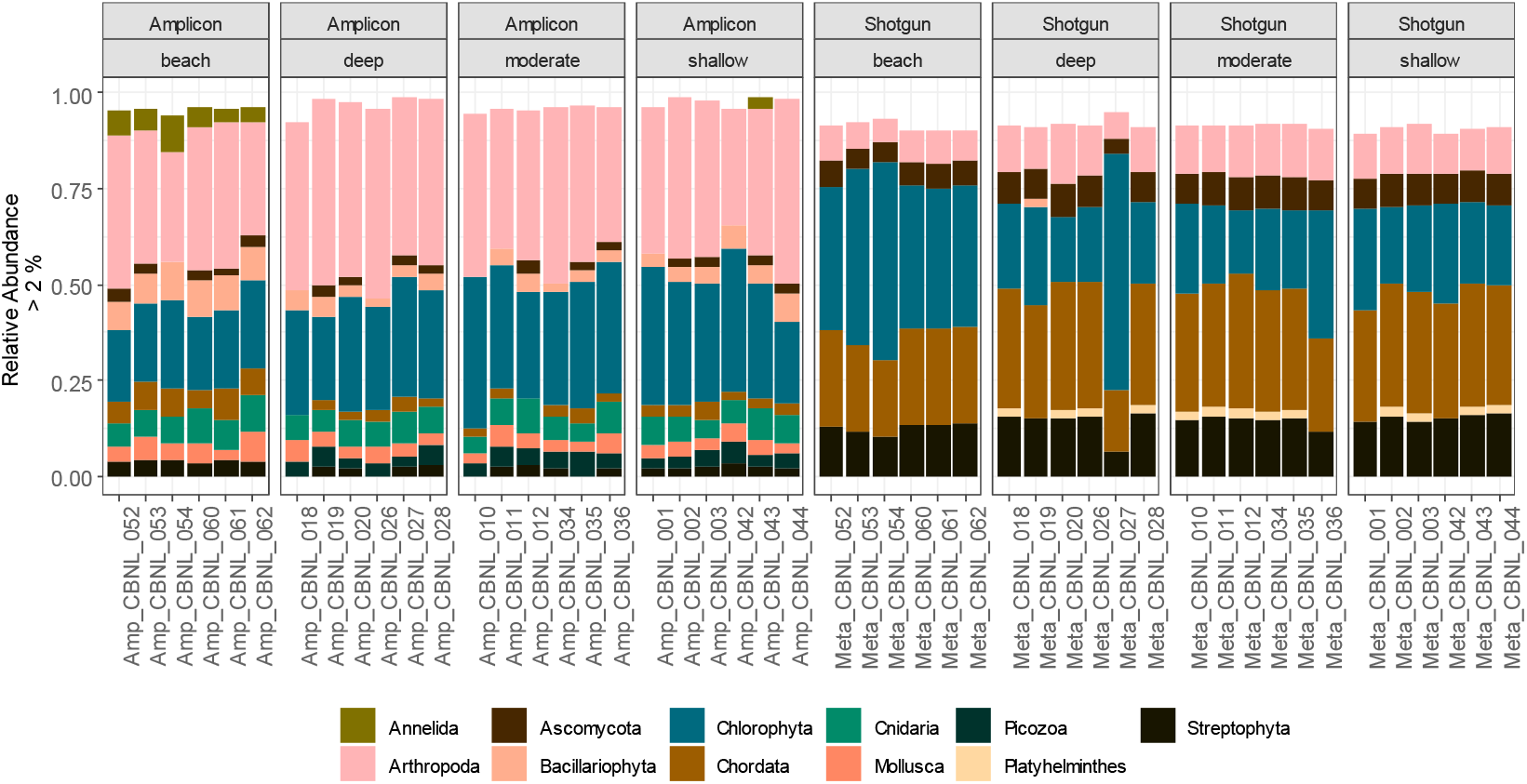
The relative proportions of identified taxa are very different between the metabarcoding experiments (left) vs the metagenomics experiment (right), although both have excellent internal consistency. Only phyla exceeding 2% are shown which is why the bars aren’t a uniform height.

**Figure 4:**
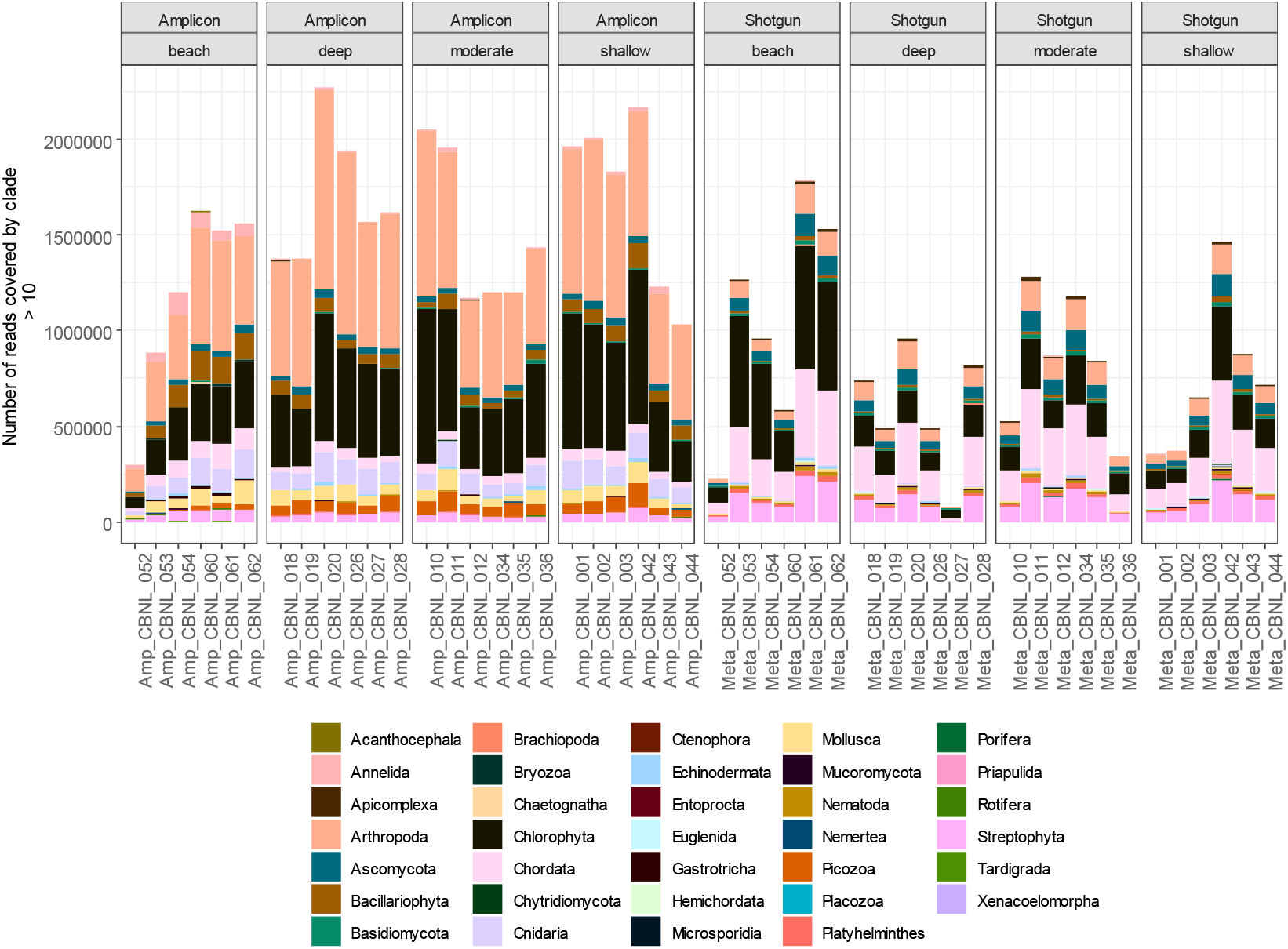
Absolute read counts for each phylum are now shown, illustrating a combination of the greater efficiency of obtaining identified taxa in metabarcoding (taller bars to the left than to the right), as well as the wide disparity in the composition of biodiversity between the two experiments.

One observation of note is that there is surprisingly little stochastic effect from one replicate to the next in the metagenomics experiments. Although this was a deep sequencing experiment, we’d still expect a very small fraction of the total eDNA to have been sequenced, and yet the taxonomic composition from one sample to the next is consistent.

### 4.3 Despite lower resolution, metagenomics data can still uncover meaningful biological patterns

Based on beta diversity, a very clear pattern emerged that showed a distinction between the water samples obtained close to shore (“beach”) and those from deeper waters (Figure 5A). Interestingly, despite the lower taxonomic resolution of the metagenomic data, the same pattern emerged in that study as well, albeit not as starkly (Figure 5B). When both metabarcoding and metagenomic data are combined, not surprisingly the primary differentiator is the methodology used, and then both methods show a difference between beach and other water samples along the secondary axis (Figure 5C). Curiously, the directionality of the difference between beach and other samples differs between the two methods, however, with beach samples higher on the y-axis than the other water samples, yet lower on the y-axis than the other water samples in the metagenomics experiments. This demonstrates that while both techniques are able to distinguish the sampling sites from each other, the exact taxa that are used to do so are completely different. In this way, a metagenomics assay can be seen as an independent confirmation of biological trends observed within metabarcoding experiments, since different taxonomic groups are likely to respond to similar ecological gradients (Ritter et al. 2019).

**Figure 5:**
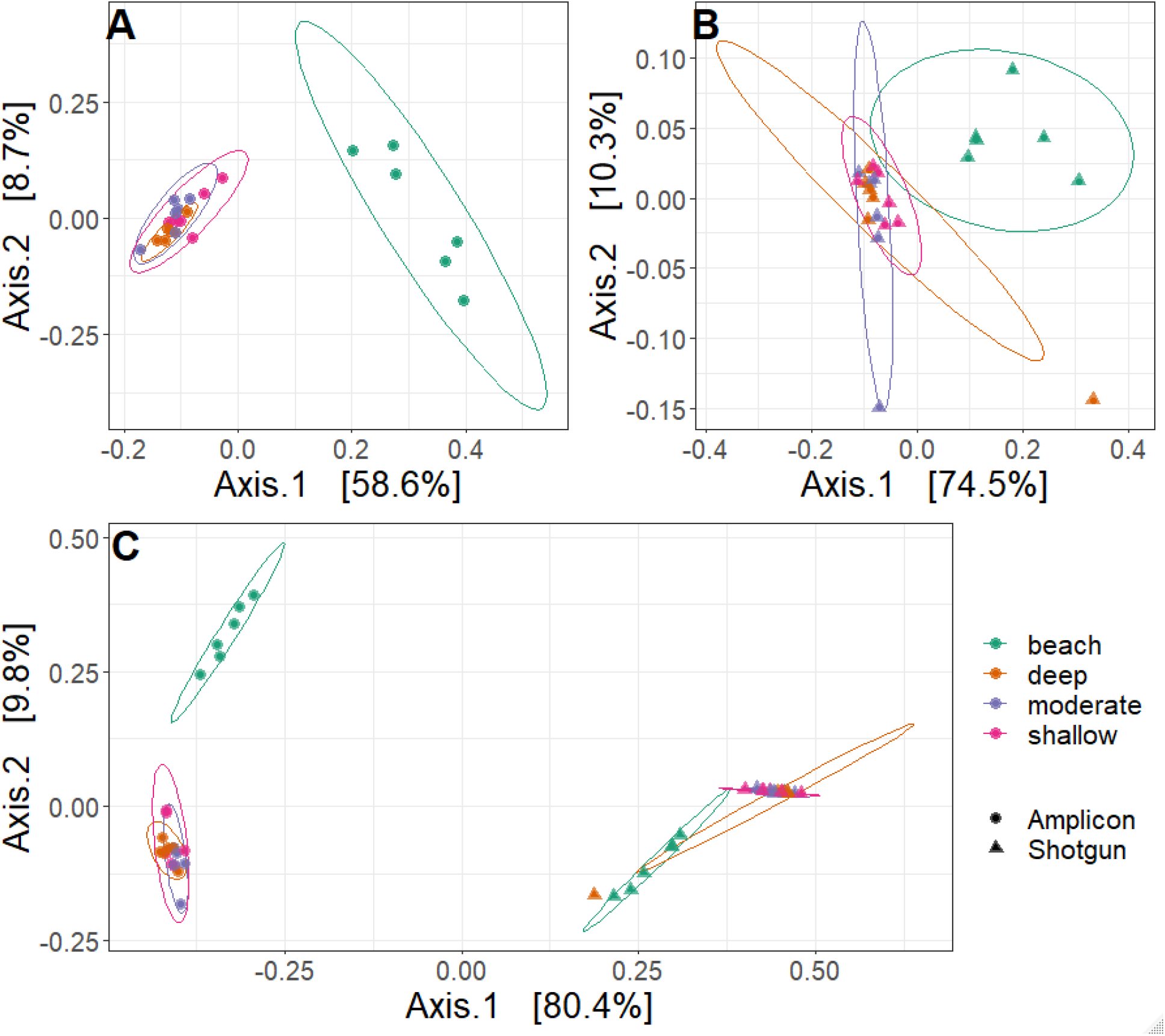
The beach samples cluster separately from the other samples in the metabarcoding data (A), and this is also true of the metabarcoding data (B), although with greater variation primarily due to two odd datapoints from deep- and moderate-depth water. When the data from both experiments are compared (C), the primary axis of variation (accounting for >80% of the variation in the data) separates the methodology (metabarcoding on the left and metagenomics on the right), while the second axis separates the beach samples from the other water samples. Interestingly, the polarity is inverted between the two experiments: the beach samples have higher values on the y-axis than the other water samples among the metabarcoding data, but this relationship is inverse among the metagenomics samples.

### 4.4 Metagenomics give us additional useful biological insights

One of the primary benefits of a metagenomics approach is that other biological information beyond simple organism identification can be obtained. Most notably, gene fragments can be identified and provide insight into the functional genomics of the environment. Here, we are talking about which genes are *present* in the environment and not which genes are *active*, which would require an RNA-seq assay (Wang, Gerstein, and Snyder 2009). Nevertheless, particularly within the microbial world, simply knowing which genes are present can give important insights into the functional diversity of the environment (Dinsdale et al. 2008). In the present analysis, we mapped shotgun sequencing reads against the CARD antimicrobial resistance gene database (Alcock et al. 2019). A principal coordinates analysis of the relative abundances of these genes reveals that the beach samples are highly distinct from the samples taken from further offshore (Figure 6A), and hierarchical clustering supports this conclusion (Figure 6B).

**Figure 6:**
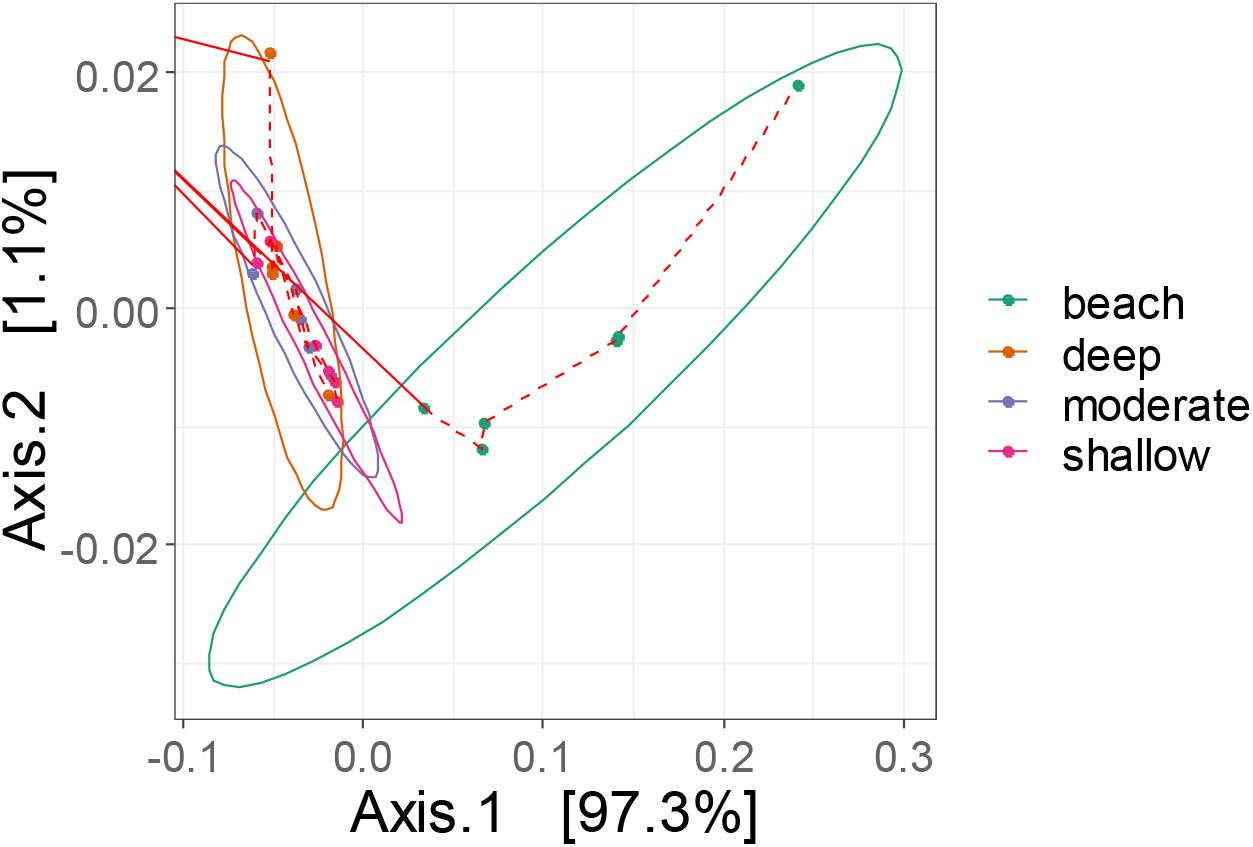
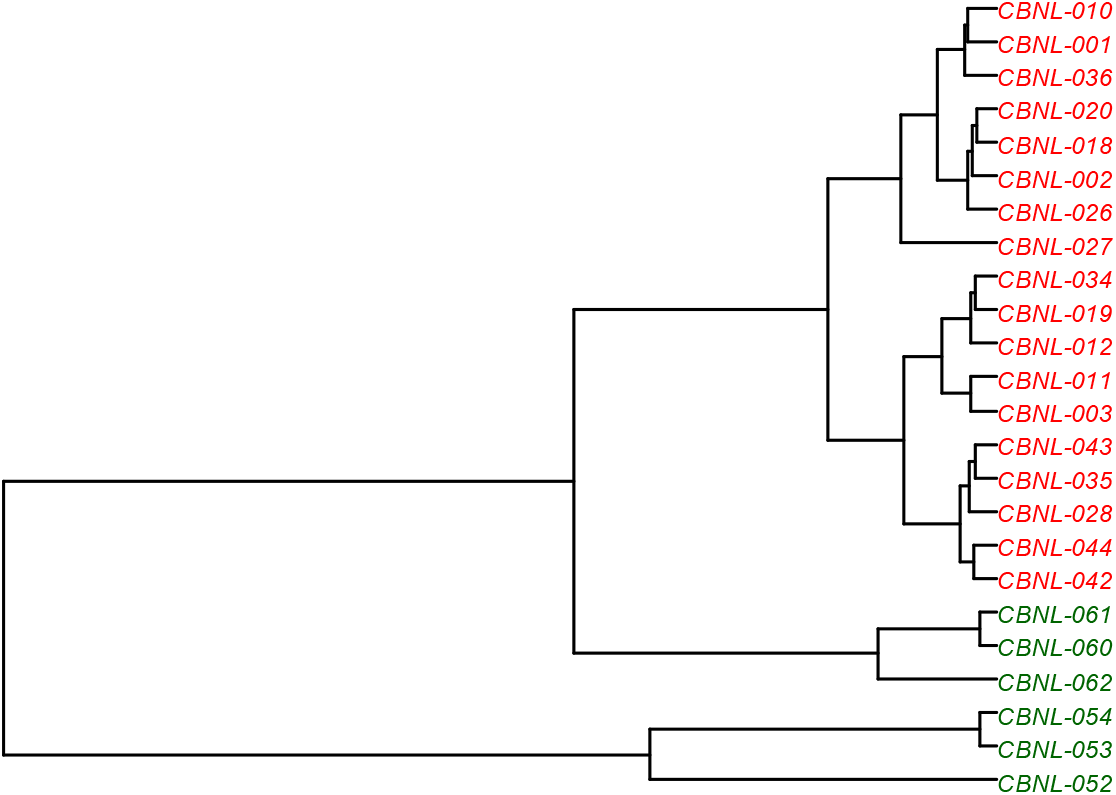
(top) PCoA based on the Bray-Curtis distances of antibiotic resistance genes. The coastal samples are clearly delineated from the non-coastal. (bottom) Hierarchical clustering of the data reveals the same pattern, with coastal (green) and non-coastal (red) samples clustering into their own clades. Interestingly, the coastal samples closest to the storm drain outlet (052, 053, and 054) are an outgroup to the rest of the samples.

To determine which gene sets were driving this difference, we plotted the relative proportions of different classes of antimicrobial resistance genes and found that the beach samples were relatively rich in antibiotic inactivation genes (Figure 7). Further analysis of these genes revealed that 24 gene families contributed to this division, 13 of which were beta-lactamases that provide resistance to broad spectrums of antimicrobials including penicillins, cephalosporins, cephamycins, and carbapenems. These genes are common within the bacterial family Enterobacteriaceae, which includes many human pathogens including *E. coli*, *Salmonella*, and *Shigella*. We turned back to the metabarcoding data to see if this family occurred in higher relative proportion in the beach samples compared to samples from further ashore, and indeed this was the case (Figure 8).

**Figure 7:**
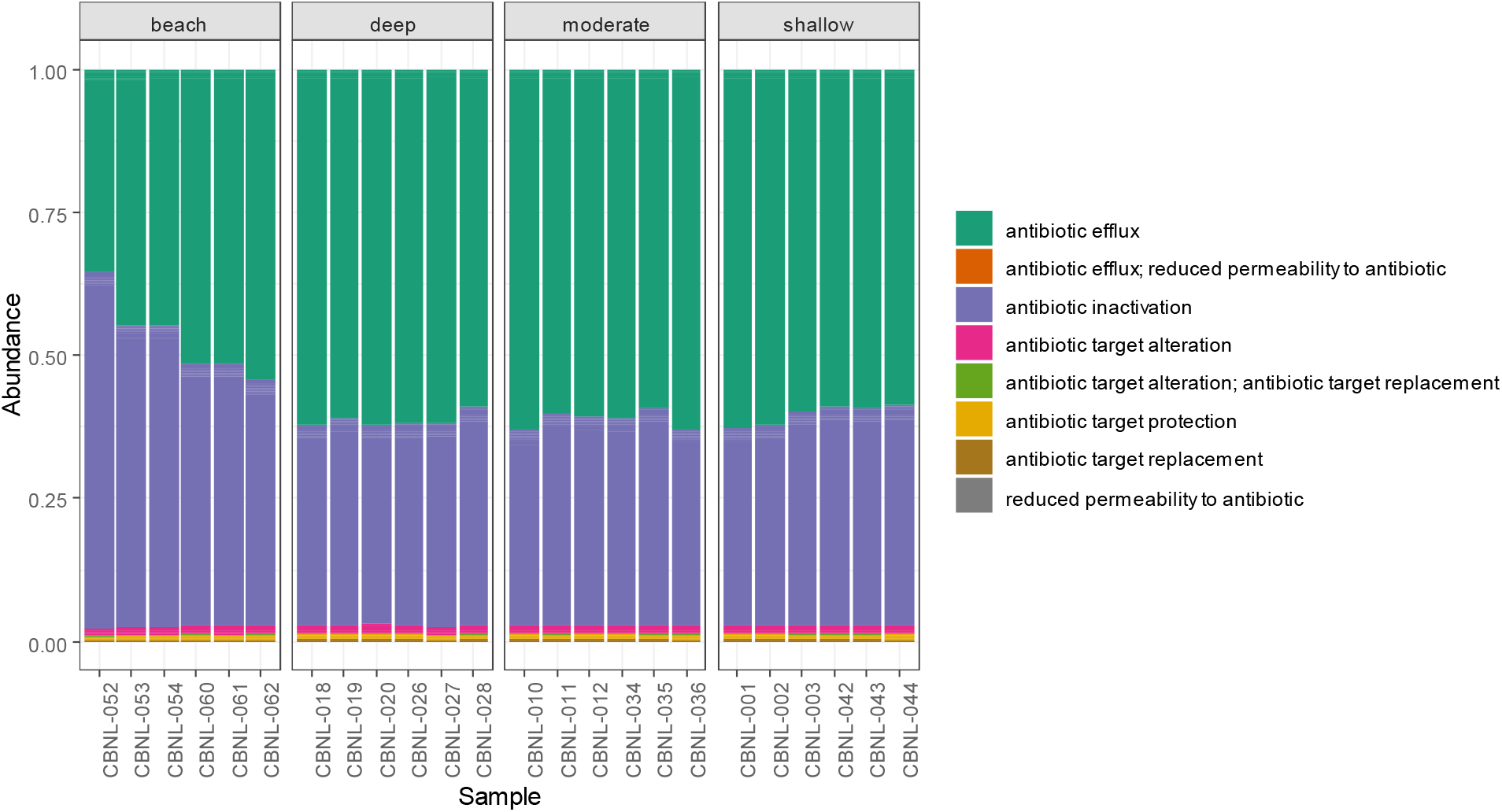
Water samples obtained close to shore have higher proportions of “antibiotic inactivation” genes present than samples taken from further offshore

**Figure 8:**
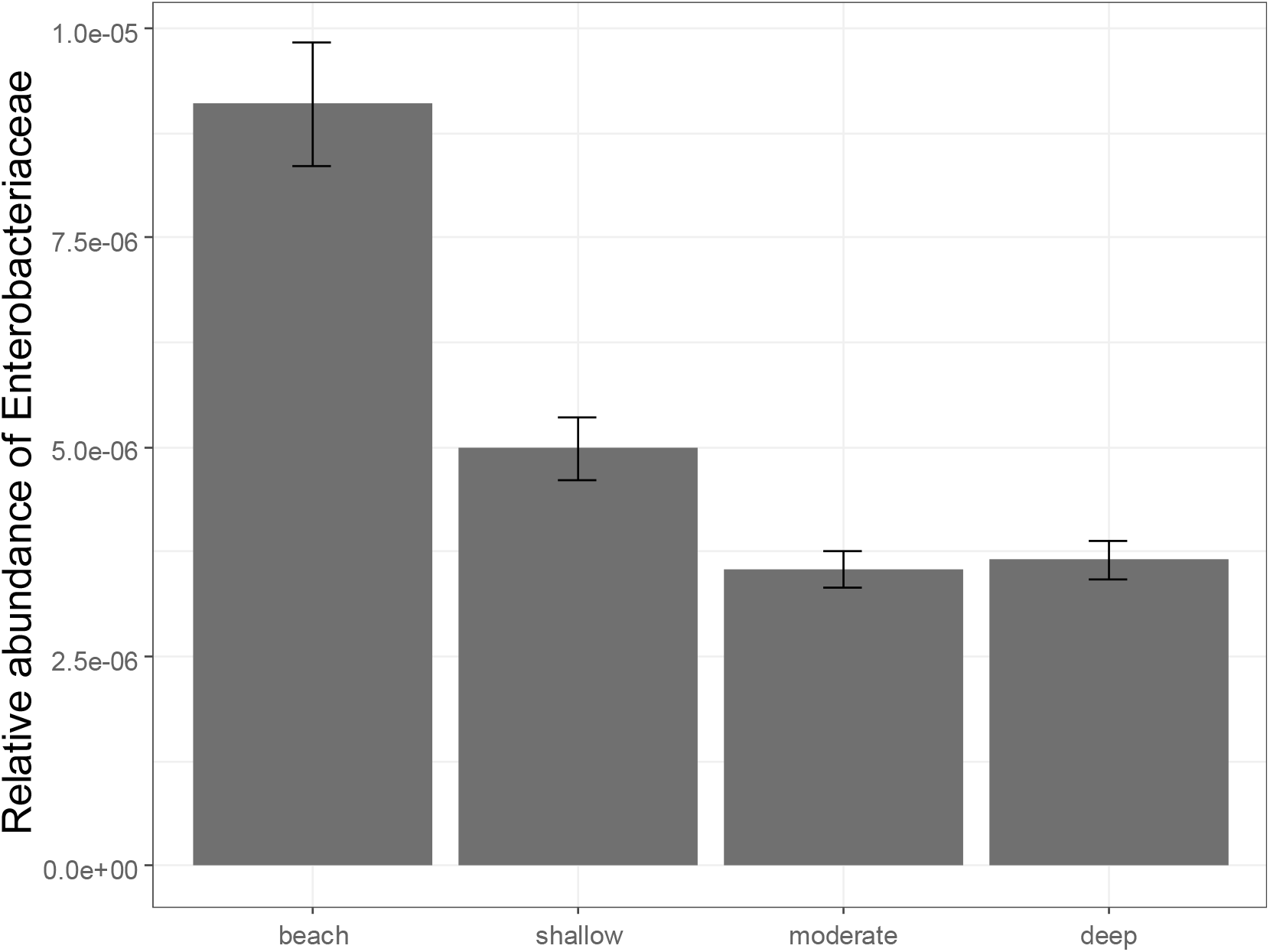
Beta-lactamase genes are known to be prevalent among the Enterobaceriacae. Here we turn back to the metabarcoding data and show that the relative proportion of this family of bacteria is indeed higher among the beach samples than in samples from deeper waters. Error bars indicate standard errors from the mean.

### 4.5 Metabarcoding experiments can be augmented with metagenomics data for “free”

It is interesting to note that on Illumina DNA sequencing platforms—the most popular instruments for metabarcoding experiments at the moment (Singer et al. 2019)—adding a metagenomics library to the flow cell can potentially be performed without adding any additional sequencing costs. This is because Illumina recommends spiking in a control library composed of PhiX genome to increase base complexity on the flow cell, with a recommended proportion of PhiX ranging from 5% to 50% (Illumina, 2017). We suggest that a shotgun environmental eDNA library could be used in place of (or, better, in combination with) PhiX to accomplish this same goal, without “wasting” space on the flow cell. Therefore, there is little downside to obtaining these useful, complementary data.

